# Speciation, sexual selection, and the macroevolution of cricket mating signals

**DOI:** 10.64898/2026.10.29.762805

**Authors:** David A. Gray, David B. Weissman, Steve Dudgeon

## Abstract

Animal mating signals can communicate both species-specific *recognition* information and male phenotypic quality *persuasion* information to females. Here we show that the recognition components of cricket song diverge early and rapidly among species pairs, especially so among those pairs with the potential for gene flow compared to allopatric pairs. Divergence in the persuasion components of song is less closely linked to speciation. Together these results suggest that the key mating signal recognition traits involved in speciation evolve in a punctuated manner, and at equilibrium are subject to stabilizing selection. Persuasion traits, on the other hand, are typically under directional sexual selection, but are not the traits closely tied to speciation. This suggests that the mechanistic link between sexual selection and speciation is ephemeral, based on rapid change in recognition traits during speciation. An important implication is that equilibrium within-population sexual selection is uncoupled from interspecific divergence during speciation. We expect that the distinction between recognition and persuasion traits, and the particular importance of divergence in recognition traits for speciation, should be a widespread phenomenon.

**Significance Statement:** Analyses of macroevolutionary patterns of cricket mating signal traits reveal mechanistically how and when rapid mating signal evolution facilitates speciation and challenges the prevailing view that traits experiencing intrapopulation sexual selection are likely to promote interspecific divergence. More than just documenting a pattern in a single system, our study provides a general framework for studying the interplay of sexual selection and speciation and for mating signal evolution generally.

## Introduction

Sexual selection was articulated by Darwin ∼150 years ago^1^, but he did not directly link it to the process of speciation, which he viewed as a consequence of an accumulation of changes driven by natural selection^2^. Mayr considered mating signals to be principally species recognition traits whose evolution fell under the umbrella of natural selection^3,4^. The more modern interest in sexual selection and speciation traces to Lande^5^ and West-Eberhard^6^, during a resurgence of interest in sexual selection generally. However, the field soon became consumed with a debate over whether mating signals are ‘arbitrary’ or ‘indicators’ of either phenotypic or genetic quality^7–9^. During this period, the role of mating signals as recognition traits was given little attention until Ryan & Rand proposed a ‘unitary’ framework in which female preference functions traverse both heterospecific and conspecific mating signal space^10^. However, the idea that separate components of mating signals serve separate functions, as either *recognition* traits or as *persuasion* traits, was not a focus of research, although it was articulated clearly by Tinbergen as early as 1953^11^, followed by Verrell^12^ and at least indirectly by others^13,14^. To this day, there has been no general reconciliation of mating signal evolution which explores recognition traits and persuasion traits separately, describes their role(s) in speciation, and describes their distinct evolutionary trajectories.

There is good reason to predict that different components of mating signals would serve as recognition traits or as persuasion traits, but that selection should operate on these functions differently, such that no single mating signal component would be well suited to both functions^15,16^. Recognition signal traits should be reliably and unambiguously associated with conspecifics, such that signal recognition promotes conspecific mating, without implying “species recognition” per se, see^17^; they should be subject to stabilizing selection^12,18,19^, and so should be stereotyped with minimal intra-population variation^15^, and relatively buffered against environmental determinants of trait expression^20^. On the other hand, attractive persuasion traits are subject to directional sexual selection^21^. Furthermore, because persistent directional sexual selection on attractive traits can promote them to develop condition dependent expression^22–25^, we expect persuasion traits may sometimes become indicators of phenotypic quality, and be phenotypically plastic and variable in all but the most benign developmental environments^26^. The development of condition dependent expression could occur regardless of whether the attractiveness of those traits was initially based on pre-existing sensory bias, or on either arbitrary ‘Fisherian’ or ‘good genes’ coevolution of traits and preferences^27^.

We also expect different macroevolutionary dynamics to be associated with recognition versus persuasion traits. Species-specific recognition traits should change during speciation, such that they remain reliably associated with conspecific identity^28^; this is especially true in sympatry or parapatry where the potential for gene flow remains. Persuasion traits, in particular condition dependent indicators, do not need to diverge at speciation; in fact, it is reasonable to expect that indicators of general condition which are attractive in one taxon would also be good indicators in another. For example, conserved mating preferences favor larger body size in many species^29–31^. These contrasting design features set up interesting macroevolutionary dynamics: recognition traits should be subject to stabilizing selection, but change at speciation, whereas persuasion traits should be subject to directional selection and yet not necessarily change at speciation^28,32^.

Testing these ideas is problematic in many taxa because mating signal components often are not clearly separable into *a priori* recognition and persuasion components (but see Discussion, below). Field crickets (Orthoptera, Gryllinae, *Gryllus* and related genera) are a clear exception to this rule: as early as 1977, Popov & Shuvalov^14^ distinguished between necessary and sufficient signal recognition traits, and signal traits which increased ‘motivation’ given the recognition of other traits. Since then, neurobiologists have been able to describe mechanistically how male cricket acoustic mating signals are processed by female cricket sensory and neural systems^33–37^, even down to the level of an individual identified neural circuit^38,39^. This body of work suggests that song fundamental frequency (FREQ), pulse rate (PRATE), and/or pulse duty cycle (PDC) function as recognition traits, whereas the overall quantity of acoustic stimulus, determined by numbers of pulses per chirp or trill (PPC), chirp or trill repetition rate (CRATE), and chirp or trill duty cycle (CDC), functions to increase the attractiveness of male song, contingent upon recognition. Gray recently showed that these mating signal traits conform to predictions based on function^15,16^: FREQ, PRATE, and PDC (i) have low intra-specific coefficients of variation, i.e. are stereotyped, (ii) are indeed subject to stabilizing selection based on female cricket response profiles, and (iii) show low phylogenetic signal, potentially indicative of rapid, but arbitrary, changes during speciation. In contrast, the persuasion traits PPC, CRATE, and CDC show high levels of intraspecific variation, are subject to directional female preferences, and show modest levels of phylogenetic signal^16^; furthermore, in 10 species of *Gryllus* crickets, Serras and Gray^20^ found that persuasion traits are often indicators of male phenotypic condition. In crickets, the recognition traits are processed by the sensory-neural system first, and persuasion trait stimulus value is processed secondarily, contingent on the signal having species-specific recognition trait properties^33,34^.

This background now allows us in this paper to compare the relative rates of evolutionary change in recognition versus persuasion traits across an entire clade of closely related cricket species (Fig. 1). We predict that recognition traits should diverge early, especially among taxa with the potential for gene flow (sympatric and parapatric taxa, see below); persuasion traits should evolve more gradually. We test these ideas using a modification of the framework outlined by Coyne and Orr^40,41^: regressing pairwise divergence in mating signals against pairwise genetic distances among species. To address the difficulty of statistical inference with phylogenetically constrained pairwise data^42^, we adopt a multi-tiered analytical approach. First, we use Bayesian regression analyses to model pairwise song divergence as a function of pairwise genetic divergence. The Bayesian approach avoids assumptions underlying frequentist statistical inference violated by non-independence of phylogenetically derived pairwise data. Instead, we use Bayesian effect sizes to contrast recognition and persuasion song traits, and ask how the potential for gene flow affects those relationships. We bolster this analysis with Multiple Regression on distance Matrices^43,44^, determining statistical significance via permutation and via comparison to a simulated null distribution of contrasts expected under Brownian Motion evolution (BM) given the phylogeny. These complementary approaches allow us to test two key predictions: (i) recognition traits are predicted to have higher intercepts and lower slopes than persuasion traits, i.e. initially evolve faster and more closely associated with speciation than persuasion traits, and (ii) this effect should be greater if gene flow is possible (i.e., in sympatry or parapatry) than in allopatry.

**Figure 1.**
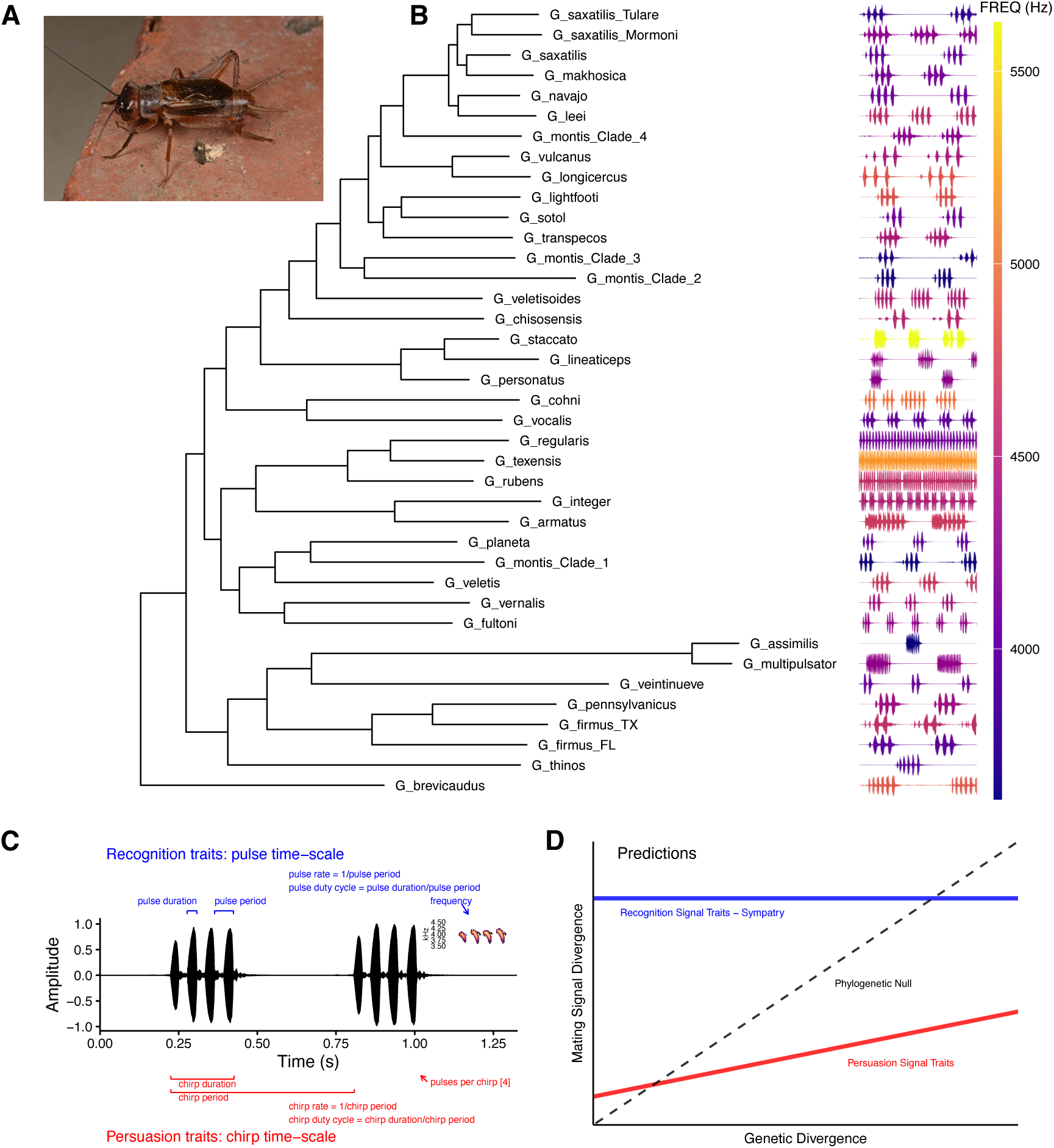
Song diversification in a clade of *Gryllus* field crickets. **A.** An adult male field cricket, *Gryllus navajo*; photo J. N. Hogue, used with permission. **B.** Phylogeny of acoustic *Gryllus* in the United States of America based on RAxML analysis of 563 loci^45^. The right side of the panel shows schematic waveform representations of 1 s of calling song per species, color coded by dominant frequency; two non-calling species (*G. ovisopis* and *G. cayensis*) were trimmed from the tree. **C.** Waveform representation of 1.25 s of song illustrating measurement and calculation of recognition song traits (pulse time scale, above in blue) and persuasion song traits (chirp time scale, below in red); inset shows frequency (kHz) of a 4-pulse chirp. **D.** Predicted evolutionary trajectories of divergence in mating signals as a function of genetic divergence among species: recognition song traits (blue) are predicted to diverge early, especially when the potential for gene flow exists, whereas persuasion song traits (red) may lag; black line shows null expectation from phylogenetic hypothesis of Brownian Motion.

## Results and Discussion

Bayesian regression analyses strongly support faster initial divergence in recognition song traits than in persuasion song traits, especially when gene flow is possible (Fig. 2 and Supplementary Materials Table S1). Recognition traits have higher intercept when in sympatry or parapatry than when allopatric, whereas the opposite is the case for persuasion traits (intercepts: Fig. 2A and 2C). Subsequent evolutionary divergence (slopes: Figs. 2B and 2D) shows the opposite patterns, with lower slope for recognition traits among species pairs when in sympatry or parapatry than when in allopatry, and accelerated slope for persuasion traits when in sympatry or parapatry. Thus, it appears that recognition trait divergence is associated with the early stages of species divergence. This interpretation is further supported by matrix regression models (MRM) in comparison with simulated phylogenetic null distributions (Fig. 3). MRM models explained significant variation in song divergence (9999 permutations, p < 0.0001 for both recognition and persuasion models). To contrast recognition and persuasion traits and control for phylogeny, we compared the observed contrasts in MRM intercepts and slopes to the distribution of contrasts expected via BM phylogenetic simulation. The observed difference in intercepts significantly differed from null expectation (p < 0.02, Fig. 3B); the observed difference in slopes was similarly unlikely to result from phylogeny alone (p < 0.02, Fig. 3B).

**Figure 2.**
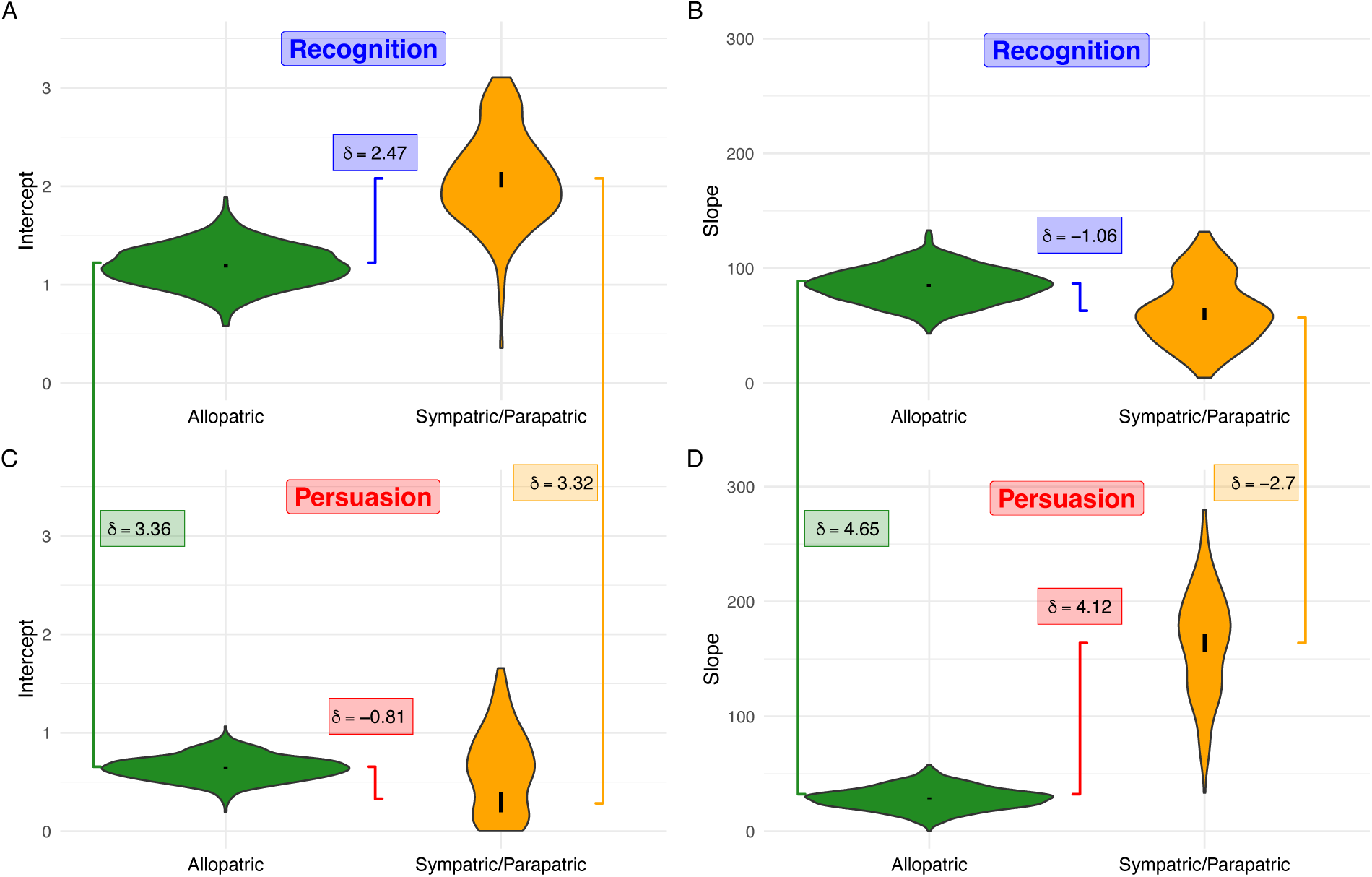
Recognition traits have higher intercepts than persuasion traits, and this effect is amplified for recognition traits in species pairs with the potential for gene flow as compared to allopatric pairs. The figure shows distributions, credible intervals, and effect size comparisons for Bayesian estimates of intercepts and slopes. Violin plots show the full posterior distributions, whereas black lines within show the 95% high density credible interval of the mean location parameters. Effect sizes of key contrasts are shown as δ values with color coded lines connecting compared means. Fig. 2A and 2C compare intercepts of recognition (2A) or persuasion (2C) trait divergence when regressed onto genetic divergence; Fig. 2B and 2D compare slopes in a similar manner, and show a lower slope for recognition trait divergence among species pairs in sympatry or parapatry than for species pairs in allopatry. Slopes for persuasion trait divergence are lower than for recognition trait divergence when in allopatry, but higher when in sympatry or parapatry.

**Figure 3.**
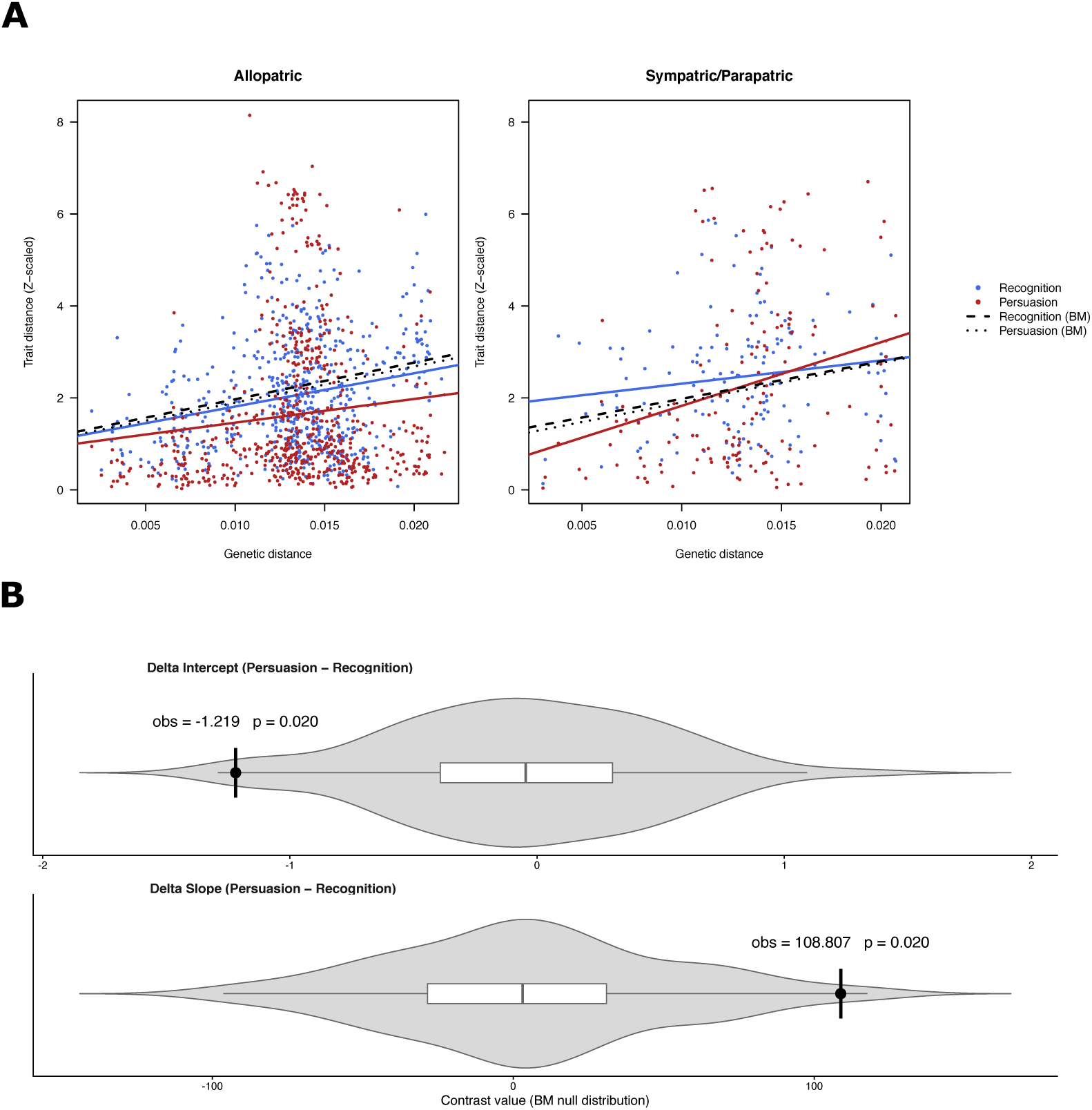
Differential trait divergence exceeds that expected by phylogeny alone. Comparison of slopes and intercepts based on MRM regression. Recognition traits show higher levels of initial divergence than do persuasion traits, especially in sympatry or parapatry when gene flow is possible (Fig. 3A). The expected effect of phylogeny is shown (black lines) based on 100 replicate Brownian Motion simulations of trait data evolution given the phylogeny. Fig. 3B shows the observed contrasts between persuasion and recognition trait intercepts and slopes in comparison to null distributions of contrast values expected based upon those phylogenetic simulations.

These results support the predictions of faster initial divergence in recognition traits than in persuasion traits, and an amplification of that effect in sympatry and parapatry where gene flow is possible, at least in the clade of acoustic *Gryllus* found in the USA. The results also underscore the importance of distinguishing between recognition and persuasion functions of different components of mating signals. That recognition traits have higher intercepts than do persuasion traits indicates faster initial divergence closely associated with speciation; that they have lower slopes indicates a subsequent reduction in the rate of divergence consistent with stabilizing selection on recognition traits at equilibrium. This suggests a ‘speciational’ model of macroevolutionary change in recognition traits. Persuasion traits, on the other hand, appear to evolve more gradually. To further explore this, we modelled trait evolution using *motmot* with both ML and Bayesian estimation of Pagel’s kappa^46^. Kappa ranges from 0 to 1, where values closer to 0 support a speciational mode, and values closer to 1 support a gradual BM mode of evolution. For ML, we compared kappa and BM models with likelihood ratio tests and with ΔAICc. Four song traits showed significant superiority of the kappa model over the BM model (Table 1): FREQ, PRATE, PPC and CDC; of the two remaining traits, PDC did not show clear evidence for either model, and CRATE appears to evolve via a gradual BM model. Thus, despite evidence of speciational change in multiple song traits, there was no clear indication that recognition traits have specially enhanced speciational change relative to persuasion traits. We interpret this result to reflect two features of cricket song evolution. First, significant phenotypic and genetic correlations among cricket song traits^47–50^ suggest that persuasion traits may change during speciation as a correlated response to change in recognition traits; therefore, both recognition and persuasion traits might be expected to show some signal of speciational effects. Second, to maintain conspecific mating signal recognition, not all of the three recognition traits need change at each node; change in any one of the recognition traits during speciation could enforce assortative mating, thus the signal of speciational change in individual recognition traits might be expected to be weakened. Evidence for this idea comes from detailed song divergence data from related sister taxa. For example, *G. personatus*, *G. staccato*, and *G. lineaticeps* form a species triplet, and it appears that ancestral *G. personatus* and the ancestor of the *G. staccato* + *G. lineaticeps* lineage diverged in frequency, but not pulse rate, and that subsequently *G. staccato* and *G. lineaticeps* diverged in pulse rate but not frequency^51^, with divergent female preferences to match^52^. Uncoupled changes among multiple recognition traits should tend to diminish the signal of speciational change in each trait individually. Speciational change in mating signals has been noted previously, particularly in insects^32,53–57^, but also in some vertebrates^58,59^.

**Table 1.** Maximum likelihood and Bayesian estimates of Pagel’s kappa, with upper and lower 95% confidence limits/credible intervals (LCI, UCI). For ML, kappa and BM models were compared via likelihood ratio tests (LRT) and ΔAICc.

|  | Maximum Likelihood |  |  |  |  | Bayesian |  |  |
|---|---|---|---|---|---|---|---|---|
| Trait | Kappa | LCI | UCI | p-value LRT | ∆AICc | Kappa | LCI | UCI |
| FREQ | 0.42 | 1.0E-08 | 0.93 | 0.025 | 2.64 | 0.48 | 0.13 | 0.85 |
| PRATE | 0.28 | 1.0E-08 | 0.88 | 0.021 | 2.99 | 0.35 | 0.04 | 0.73 |
| PDC | 0.42 | 1.0E-08 | 1 | 0.123 | 0.03 | 0.45 | 0.07 | 0.89 |
| PPC | 0.28 | 1.0E-08 | 0.88 | 0.023 | 2.8 | 0.38 | 0.07 | 0.78 |
| CRATE | 1 | 0.18 | 1 | 1 | -2.35 | 0.66 | 0.27 | 0.97 |
| CDC | 0.06 | 1.0E-08 | 0.8 | 0.009 | 4.48 | 0.34 | 0.04 | 0.75 |

Appreciating the distinction between recognition and persuasion functions of different components of mating signals helps clarify the expected relationship between sexual selection and speciation. For several decades, a variety of authors have suggested that sexual selection may play an important role as an accelerant of divergence in prezygotic reproductive isolation, reviewed in^60–63^. In some models, authors envision divergent mate preferences alone directly causing speciation^e.g., 64^, whereas in others sexual selection plays a supporting role promoting assortative mating among ecologically divergent populations^e.g., 65^. In our view, these kinds of models are conceptually most applicable to recognition traits. However, given that recognition traits should have low variation and be under stabilizing selection at equilibrium, recognition traits may not be readily identified by empirical researchers as being ‘sexually selected’ – at equilibrium, intraspecific variation in recognition traits should be low and minimally associated with variation in mating success within a population. Persuasion mating signal traits are much more easily identified as being under sexual selection given that they are variable, subject to directional mating preferences, and may often be condition dependent. Thus, despite the great interest in sexual selection and speciation, we should not expect that most mating signal traits which researchers readily identify as ‘sexually selected’ would be especially associated with speciation. This counter-intuitive result has in fact been supported by several recent studies specifically designed to test if those traits under strong within-population sexual selection are the same traits which show important divergence among populations or species. For example, an elegant study on wolf-spider communication signals showed that male mating traits currently subject to sexual selection are not those most divergent among species^66^. A similar

study in *Teleogryllus* field crickets, but at the genomic level, found that the genes associated with intraspecific mating traits were not the genes associated with interspecific divergence^67^. Instead, we should expect the mechanistic link between sexual selection and speciation to be ephemeral, based on rapid change in recognition traits during speciation. When comparing groups of species, we should therefore expect that the degree of divergence in recognition traits and preferences would be associated with speciation, rather than expecting that the strength of sexual selection *per se* would be associated with speciation; such a pattern has in fact been found in a variety of taxa^68–70^.

How general are these results, and how can researchers distinguish between recognition and persuasion traits in other taxa? We expect that the distinction between recognition and persuasion traits, and the particular importance for speciation of divergence in recognition traits, should be a widespread phenomenon likely to become more apparent with careful study of potential trait functions. We are not aware of previous work which has both distinguished between recognition and persuasion signal functions and shown different evolutionary trajectories for those different trait functions. However, a large number of studies have identified signal properties consistent with at least some of these ideas. For example, orchid bee perfumes^71^, *Enchenopa* treehopper vibrations^72,73^, Gomphocerinae grasshopper calls^74^, *Tetragnatha* spider silks^75^, *Teleogryllus*^76,77^ and *Laupala*^78^ cricket songs, claw displays of *Uca* fiddler crabs^79–81^, and bovid and cervid display complexity^82^ all share similarities with this work. On the other hand, studies in classical lekking bird systems examining whether suites of complex mating signal traits show correlated or independent evolution have given conflicting results, e.g. contrast manakins^83^ and birds-of-paradise^84^; we would expect that on-lek displays may often include a mix of recognition and persuasion traits, but we would also expect that calls which function from a distance to advertise the location and/or activity of the lek would primarily be recognition displays. In other words, identification of recognition and persuasion traits can be complicated, but generally we should expect that recognition traits (i) are displayed early in mating sequences and/or given sensory-neural priority, (ii) are stereotyped, with low intraspecific variation and minimal phenotypic plasticity, (iii) at equilibrium are under stabilizing selection from signal receivers.

## Methods

### General

Analyses were conducted in R 4.2.1.

### Taxa, phylogeny and evolutionary divergence

We analyze divergence in the long-distance mating signals of all currently recognized lineages of *Gryllus* crickets in North America north of Mexico which produce calling song: 33 named species plus 6 genetically distinct but otherwise cryptic lineages which are currently not named^85^; for brevity, we refer to the set of 39 collectively as ‘species.’ Two named species were excluded (*G. ovisopis* and *G. cayensis*) as they do not produce calling song^86–88^. Phylogenetic relationships were as described in Gray et al.^45^ based on RAxML analysis of 563 concatenated loci (492,531 bp) from anchored hybrid enrichment. We calculated the matrix of patristic pairwise evolutionary distances, i.e. sum of phylogenetic branch lengths, between pairs of taxa using the cophenetic.phylo function in *ape*.

### Mating signals and song divergence

For each lineage, we analyzed the calls of five wild-caught males recorded in captivity at ∼25 °C, for details see^15,16^. Briefly, using Audacity software, we measured five exemplars of each trait for each individual (see Fig. 1C), averaged the values within individuals and then averaged the five individuals within each species to obtain species-typical values for frequency, pulse rate, pulse duty cycle, number of pulses per chirp, chirp rate, and chirp duty cycle. We then used the dist function (method = “euclidean”) in the base *stats* package to create pairwise song distance matrices for recognition and persuasion traits calculated separately on Z transformed song data.

### Potential for gene flow

We coded all pairwise species comparisons as either ‘Allopatric’, ‘Parapatric’ or ‘Sympatric’ based on known collection localities^85^. The categories were intended to capture the potential for gene flow in the absence of intrinsic mating barriers: allopatric = geographically separate, no potential gene flow; parapatric = geographically and/or ecologically adjacent, some gene flow possible at range edges and/or ecotones; sympatric = extensive geographic overlap, gene flow likely unless prevented by intrinsic barriers.

### Bayesian regression models

Divergence of cricket songs between groups differing in potential gene flow and recognition or persuasion components of song was estimated by regressing pairwise song distances on evolutionary distances among species pairs comprising each group. Rates and magnitudes of divergence of song with evolutionary distance were estimated from slopes and intercepts of each group. The complete set of 741 interspecific pairwise distances in a matrix of the 39 species consisted of 594 allopatric, 77 parapatric, and 70 sympatric species pairs. Given the much smaller sample sizes for the latter two and our primary interest in comparing divergences in song between species with and without the potential for gene flow, we pooled parapatric with sympatric species pair groups to have n=147 pairs of species with potential gene flow for regression analyses. For each potential gene flow group, we analyzed song divergence separately for recognition or persuasion traits (i.e., 4 groups in total; allopatric-recognition, allopatric-persuasion, gene flow-recognition, gene flow-persuasion).

We constructed Bayesian linear regression models using jags for MCMC sampling in R^89^. Models were robust to data that were not normally distributed owing to heavy-tailed distributions by modeling the data as t-distributed with normality parameter, ν, and estimating the standard deviation of song divergence as well as the intercept and slope. Regressions were done on standardized (Z-transformation) song and branchlength distance data and converted back to conventional parameters for interpretation and presentation.

All parameters were modeled using vague priors on the standardized scale. Regression coefficients were modeled as normally distributed with mean equal to 0 and standard deviation equal to 10. Standard deviations of song distance were modeled as uniformly distributed between 10^-3^ and 10^3^. The normality parameter, ν, of the t-distribution assumed an exponential distribution with mean (parameter λ) equal to 30 which provides equal opportunity to values greater, or less, than 30 where most of the variation in the shape of *t* occurs as it converges towards the normal when ν=30^89^.

Posterior distributions for each random variable were generated from Markov Chain Monte Carlo (MCMC) samples of four chains run in parallel. Each chain consisted of 500 adaptation steps to tune parameters, plus 1000 burn-in steps followed by 5,000 saved steps that were not thinned to yield 20,000 saved steps across the four chains. Diagnostics of MCMC chains, including trace plots, the Gelman-Rubin statistic, and autocorrelation were examined to confirm convergence of values among chains for each parameter. The median effective sample sizes (ESS) for regression parameters, β0, β1, and σ each exceeded 11,400 for all 4 groups except for σ among allopatric-recognition and allopatric-persuasion groups that were 5455 and 4735, respectively. The 95% high-density (credible) intervals of the posterior distributions of regression parameters were compared relative to 0 to infer their magnitude and variability, and slopes (β1) were also contrasted against a region of practical equivalence (ROPE) with respect to 0 of -0.5 to +0.5.

Two-group comparisons of the posterior distributions of slopes or intercepts were made between either allopatric versus gene flow groups (separately for recognition and persuasion song trait groups), or between recognition versus persuasion song trait groups (separately for allopatric or gene flow groups). For these two-group tests, 594 observations from the allopatric group and 147 observations from the gene flow group were drawn randomly from their respective posterior distributions of regression parameters for both recognition and persuasion traits to generate new distributions which reflected the intensity of actual sampling in each comparison. Data for these two-group tests were modeled as t-distributed with normality parameter, ν, with separate estimates of variance for each group. Priors for model parameters were, again, vague. Population parameters, μ, were modeled as normally distributed with mean equal to the mean of the sampled data set and very low precision (the reciprocal of the square of 100 times the standard deviation). Scale parameters were modeled as uniformly distributed between the standard deviation times 10^-3^ and 10^3^. The normality parameter assumed an exponential distribution with λ= 30 as above.

Posterior distributions for each sample as well as the posterior distribution of the difference between samples were generated from MCMC samples of four chains run in parallel. Each chain consisted of 500 adaptation steps, 1000 burn-in steps followed by 12,500 unthinned saved steps (50,000 saved steps across the four chains). Diagnostics of MCMC chains, including trace plots, the Gelman-Rubin statistic, and autocorrelation were examined to confirm convergence of values among chains for each parameter. The 95% high-density intervals of the posterior distributions of each parameter were compared to one another to infer the magnitude of their difference and also contrasted against a difference of 0 between posterior distributions and a ROPE of -0.1 to +0.1 with respect to the difference in location between distributions.

### Phylogenetic null and Matrix regression models

In order to generate a set of null datasets which preserve the observed scale and phylogenetic covariance structure of the data, we simulated neutral song evolution on the tree using a Brownian motion (BM) model implemented via rTraitCont in *ape*^90^. For each trait we generated 100 replicate datasets. Frequency, pulse rate, pulses per chirp, and chirp rate, which are biologically constrained to positive values, were each simulated on the log scale ensuring positive values after back transformation; similarly, pulse and chirp duty cycles, biologically constrained to values between 0 and 1, were simulated on the logit scale to preserve those constraints in the simulated data after back transformation. For each trait, simulations started with the transformed observed mean at the root and with the Brownian Motion variance parameter set such that the expected tip variance would match the observed variance for each trait. This was done by setting the variance parameter equal to the observed trait variance divided by the mean of the diagonal elements of the phylogenetic variance-covariance matrix computed by vcv.phylo function in *ape*^90^. For each of the 100 simulated data sets, we then calculated a simulated song distance matrix as described above under ***Mating signals and song divergence***.

For both the observed and 100 simulated data sets, we then used Multiple Regression on distance Matrices with the MRM function in *ecodist*^91^ to model song divergence **Y** on genetic distance **X** + potential for gene flow **P** + the **XP** interaction where **XP** was calculated externally to the model as **X*P**. Statistical significance of the overall model was determined relative to 9999 permutations. To further control for phylogeny, statistical significance of observed trait contrasts was determined via comparison to contrasts from the simulated 100 data sets.

### Song Trait Evolution

Models of song trait evolution on phylogenetic trees were run using *motmot*^92^. Each of the 6 song traits (3 recognition – frequency, pulse rate, pulse duty cycle; 3 persuasion - number of pulses per chirp, chirp rate, and chirp duty cycle), were integrated with data on tree structure and plotted on a phylogeny. Models of evolution were analyzed separately for each trait using both ML and a Bayesian MCMC algorithm to estimate whether trait change occurs principally at speciation (kappa approaching 0) or gradually throughout a phylogeny (kappa approaching 1). BM and kappa ML models were compared via likelihood ratio tests and ΔAICc. Bayesian models had uniform distributions for prior estimates of parameters such that all values were equally likely. Each MCMC chain consisted of 5000 iterations, thinned every 5 yielding 1000 saved steps, of which the first 100 were burn-in, totaling 901 saved steps per chain. Four MCMC chains were run for each estimated parameter for a total of 3604 saved steps to the posterior distribution. The 95% High Density interval of each posterior distribution was compared to informative values of parameters (e.g., 0 or 1).

## Data availability

Data will be made publicly available in Dryad.

## Code availability

Code for all analyses will be publicly available in Dryad.

## Notes

https://data.mendeley.com/datasets/7tprb8zhgt/1

### Competing Interest Statement

The authors have declared no competing interest.

