## Supplemental Material Table S1 for "Speciation, sexual selection, and the macroevolution of cricket mating signals"

Table S1. Regression parameters from posterior distributions compared among potential for genetic mixing (Allopatric or Gene Flow) and type of song trait (Recognition or Persuasion). Estimates are based on random samples of 594 or 147 observations from the posterior distributions to reflect actual sampling intensities for allopatric and gene flow comparisons, respectively. HDI refers to the high density (95%) credible interval between low and high values. Contrasts calculated as Gene Flow - Allopatric or Recognition – Persuasion; Effect size calculated as the difference in location divided by the averaged standard deviation.

| **Recognition Trait Comparisons** | | | | | | |
|---|---|---|---|---|---|---|
| Parameter | Contrast | Mean | Median | Mode | HDI Low | HDI High |
| $\beta 0$ | Allopatric | 1.192 | 1.192 | 1.191 | 1.176 | 1.208 |
| | Gene Flow | 2.068 | 2.068 | 2.070 | 1.990 | 2.145 |
| | Posterior location difference | 0.876 | 0.876 | 0.877 | 0.797 | 0.955 |
| | Effect Size | 2.473 | 2.471 | 2.453 | 2.131 | 2.830 |
| $\beta 1$ | Allopatric | 85.153 | 85.153 | 85.167 | 83.981 | 86.334 |
| | Gene Flow | 60.145 | 60.145 | 60.221 | 55.099 | 65.170 |
| | Posterior location difference | -25.008 | -25.010 | -24.660 | -30.090 | -19.737 |
| | Effect Size | -1.057 | -1.057 | -1.056 | -1.294 | -0.813 |
| **Persuasion Trait Comparisons** | | | | | | |
| Parameter | Contrast | Mean | Median | Mode | HDI Low | HDI High |
| $\beta 0$ | Allopatric | 0.641 | 0.641 | 0.641 | 0.631 | 0.651 |
| | Gene Flow | 0.291 | 0.290 | 0.289 | 0.192 | 0.392 |
| | Posterior location difference | -0.350 | -0.351 | -0.355 | -0.450 | -0.248 |
| | Effect Size | -0.808 | -0.808 | -0.813 | -1.060 | -0.560 |
| $\beta 1$ | Allopatric | 28.610 | 28.611 | 28.621 | 27.845 | 29.369 |
| | Gene Flow | 164.008 | 163.993 | 163.881 | 156.419 | 171.404 |
| | Posterior location difference | 135.397 | 135.380 | 135.362 | 127.800 | 142.874 |

| | | | | | | |
|---|---|---|---|---|---|---|
| | Effect Size | 4.122 | 4.117 | 4.084 | 3.584 | 4.656 |
| **Allopatric**<br>**Posterior location differences** | | | | | | |
| Parameter | Contrast | Mean | Median | Mode | HDI Low | HDI High |
| $\beta0$ | Recognition - Persuasion | 0.551 | 0.551 | 0.551 | 0.532 | 0.570 |
| | Effect Size | 3.361 | 3.359 | 3.358 | 3.152 | 3.574 |
| $\beta1$ | Recognition - Persuasion | 56.528 | 56.530 | 56.425 | 55.117 | 57.916 |
| | Effect Size | 4.651 | 4.649 | 4.653 | 4.404 | 4.909 |
| **Gene Flow**<br>**Posterior location differences** | | | | | | |
| Parameter | Contrast | Mean | Median | Mode | HDI Low | HDI High |
| $\beta0$ | Recognition - Persuasion | 1.776 | 1.776 | 1.770 | 1.648 | 1.902 |
| | Effect Size | 3.322 | 3.318 | 3.316 | 2.939 | 3.725 |
| $\beta1$ | Recognition - Persuasion | -103.955 | -103.977 | -104.229 | -112.943 | -94.888 |
| | Effect Size | -2.703 | -2.699 | -2.683 | -3.064 | -2.351 |
